# STAN, a computational framework for inferring spatially informed transcription factor activity

**DOI:** 10.1101/2024.06.26.600782

**Authors:** Linan Zhang, April Sagan, Bin Qin, Elena Kim, Baoli Hu, Hatice Ulku Osmanbeyoglu

## Abstract

Transcription factors (TFs) drive significant cellular changes in response to environmental cues and intercellular signaling. Neighboring cells influence TF activity and, consequently, cellular fate and function. Spatial transcriptomics (ST) captures mRNA expression patterns across tissue samples, enabling characterization of the local microenvironment. However, these datasets have not been fully leveraged to systematically estimate TF activity governing cell identity. Here, we present STAN (<u>S</u>patially informed <u>T</u>ranscription factor <u>A</u>ctivity <u>N</u>etwork), a linear mixed-effects computational method that predicts spot-specific, spatially informed TF activities by integrating curated TF-target gene priors, mRNA expression, spatial coordinates, and morphological features from corresponding imaging data. We tested STAN using lymph node, breast cancer, and glioblastoma ST datasets to demonstrate its applicability by identifying TFs associated with specific cell types, spatial domains, pathological regions, and ligand‒receptor pairs. STAN augments the utility of STs to reveal the intricate interplay between TFs and spatial organization across a spectrum of cellular contexts.

## Introduction

Elaborately orchestrated transcriptional programs distinguish specialized cell types/states and define their functionality. Combinations of transcription factors (TFs) drive these programs through interactions with *cis*-regulatory elements, such as promoters and enhancers, to control cellular identity and functional state. Nearby cells are critical for determining context-specific TF activity. TFs are typically expressed at low levels and are regulated post-transcriptionally and/or post-translationally. Furthermore, cellular processes depend on the expression levels (and activities) of proteins, notably TFs, which can be distinct from those at the mRNA level.

TF activities are commonly inferred from their target gene expression. Various methods to investigate TF activity have been developed based on bulk [1–3] and single-cell omics [4] datasets. Early methods using single-cell omics relied on single-cell RNA-seq (scRNA-seq) data and/or the identification of TF motifs in annotated promoter regions [4–10]. These methods primarily depend on the coexpression of TFs and their potential target genes (e.g., the SCENIC tool [4]). Given the availability of single-cell epigenomic datasets such as those generated by single-cell sequencing assays for transposase-accessible chromatin, various TF inference methods have focused on context-specific accessible regions of chromatin that may be bound by TFs either upstream or downstream of target genes. Recently, SPaRTAN (Single-cell Proteomic and RNA-based Transcription Factor Activity Network) was developed, which leverages single-cell proteomic and corresponding scRNA-seq datasets obtained via the cellular indexing of transcriptomes and epitopes by sequencing, to link surface proteins with TFs [11]. However, these approaches overlook relationships between other signaling proteins and TFs within individual cells, as well as the influence of surrounding cells on these interactions and how cell identity is affected by the local environment.

Genome-wide spatial transcriptomics (ST) platforms such as 10x Genomics’ Visium ST, DBiTseq [12], and Slide-seq [13] enable transcriptome profiling of cells in their native context and provide information about how cell identity is influenced by locality. For example, Visium measures up to 5,000 spots, each containing up to 10 cells. ST has been used to study cancer [14–18], Alzheimer’s disease, periodontitis [19, 20], and healthy tissues [21, 22]. One advantage of ST data is the additional generated information of coregistered images, which contain both morphological and functional patterns. Multiple computational approaches have been introduced to analyze ST data, including the characterization of spatial gene expression patterns, spatially distributed differentially expressed genes, and spatial cell‒cell communication patterns [23–34]. Despite these advances, no existing methods integrate spatial and histological data with gene expression and *cis*-regulatory information to systematically estimate spot-specific TF activities.

Here, we describe a new linear mixed-effects model computational framework, STAN (<u>S</u>patially informed <u>T</u>ranscription factor <u>A</u>ctivity <u>N</u>etwork), for inferring spatially informed TF activity (**Figure 1**). This framework leverages curated TF-target gene priors, spot-specific gene expression, spatial coordinates, and morphological features extracted from corresponding hematoxylin and eosin (H&E) histological data obtained with ST. Briefly, STAN views the expression of TF-target genes as a proxy of their activities. Spatial coordinates and morphological features extracted from corresponding imaging data are used to estimate the spatially informed activity of TFs. Widely used ST platforms, such as Visium, measure the entire transcriptome at low-resolution spots, limiting their ability to detect the activity of cell type-specific TFs. Therefore, STAN incorporates cell-type deconvolution approaches (e.g., Cell2location [35], DestVI [36], Tangram [37], Stereoscope [38], RCTD [39], and others) to identify cell type-specific TFs. We have successfully applied and tested STAN using lymph node, breast cancer, and glioblastoma ST datasets to demonstrate its utility by identifying TFs associated with specific cell types, spatial domains (e.g., germinal centers [GCs]), pathological regions, and ligand‒receptor pairs.

**Figure 1:**
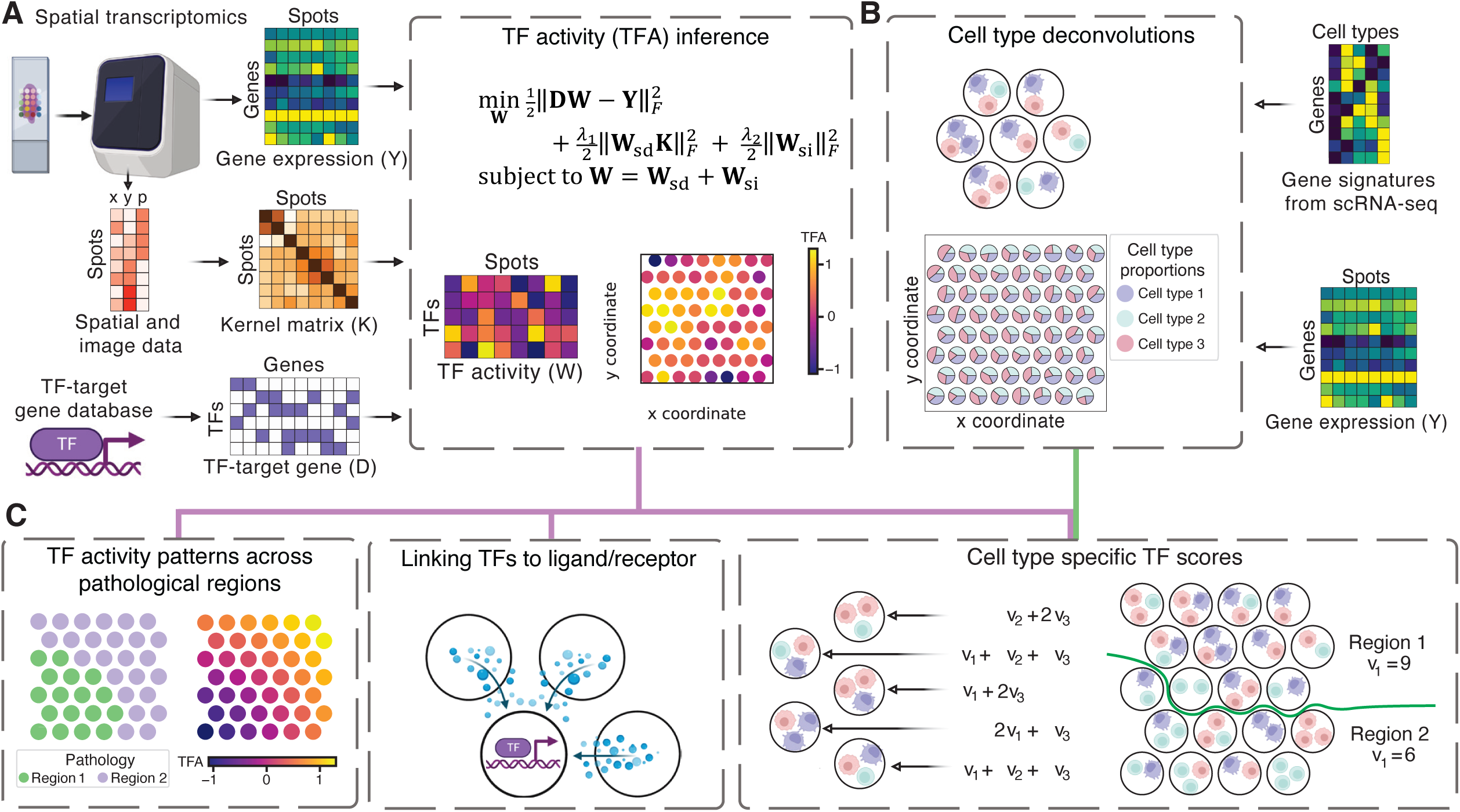
Overview of the STAN workflow. **(A)** As input, our model accepts input as a normalized gene expression matrix, spatial locations, and pixel intensity of the corresponding image for each spot, along with a binary matrix indicating the genes each TF targets as obtained via curated database TFs. Spot-specific TF activity levels are inferred by training a spatially regularized linear regression model to predict gene expression via TF-target gene relationships. **(B)** Deconvolution analysis infers spot-specific cell type/state proportions on the basis of reference scRNA-seq data (signature matrix). Downstream analysis includes **(C)** inferring cell type-specific TFs, **(D)** identifying TF activity patterns across distinct pathological regions and spatial domains, and **(E)** identifying ligands/receptors whose gene expression correlates with TF activity in neighboring spots.

## Results

### Inferring TF activity according to cell type, physical location, and ligand/receptor associations

STAN is a toolbox for analyzing spot-based ST datasets to estimate spot-specific TF activities and identify TFs associated with specific cell types, spatial domains, pathological regions, and ligands/receptors (**Figure 1**). STAN uses spot-specific morphological information, spatial coordinates, and gene expression (**Y**), together with TF-target gene priors (**D**), as inputs and then models gene expression in terms of spatially coherent TF activities (**Figure 1A**). To achieve this capability, we used a kernel regression model, where we created a spot-specific TF activity matrix, **Y,** that is decomposed into two terms: one required to follow a spatial pattern (**W**_sd_) generated via a kernel matrix (**K**) and another that is unconstrained but regularized via the *L*_2_-norm (**W**_si_) The kernel matrix enforces similarity in spot-specific TF activity in nearby physical spaces and/or morphologically. We constructed **K** via spatial proximity between spots (Euclidean distance) and morphological similarity (measured as the similarity between pixel intensities for H&E-stained images). The model then fits the expression of each gene as the sum of the activities for a set of TFs.

A key limitation of most grid-based ST profiling methods is the lack of single-cell resolution, as spatial RNA-seq measurements combine multiple cells (e.g., 1–10 cells). Hence, the inferred spot-specific activity of TFs is from multiple cells. To overcome this limitation, we used deconvolution methods to estimate cell type proportions for each spot via the computational integration of spatial RNA-seq with the reference transcriptome signatures of cell types obtained from scRNA-seq profiles (**Figure 1B**). We then performed association analysis between TFs and cell types via linear regression to predict the activity of each TF individually from the cell type proportions via deconvolution methods (**Figure 1C**). We also performed downstream analysis to identify TFs whose activities differ across pathological regions and spatial domains (**Figure 1D**) and to identify ligands/receptors whose gene expression is associated with TFs in neighboring locations (**Figure 1E**).

### Identifying GCs and cell type-specific TFs in the human lymph node

We first trained STAN using a publicly available Visium ST dataset from a human lymph node (**Supplementary Figure 1A**). For statistical evaluation, we computed the PCC between the predicted and measured gene expression profiles at held-out spots via 10-fold cross-validation. We obtained significantly better performance than the baseline model without spatial/morphological information based on ridge regression (*P* < 10^-16^, one-sided Wilcoxon rank-sum test; **Figure 2A**). In particular, the increased benefit of spatial information and prediction performance was inversely correlated with the total UMI count per spot (PCC = -0.2097), suggesting that our method of pooling information from neighboring spots was more effective than utilizing spots with low UMI counts.

**Figure 2:**
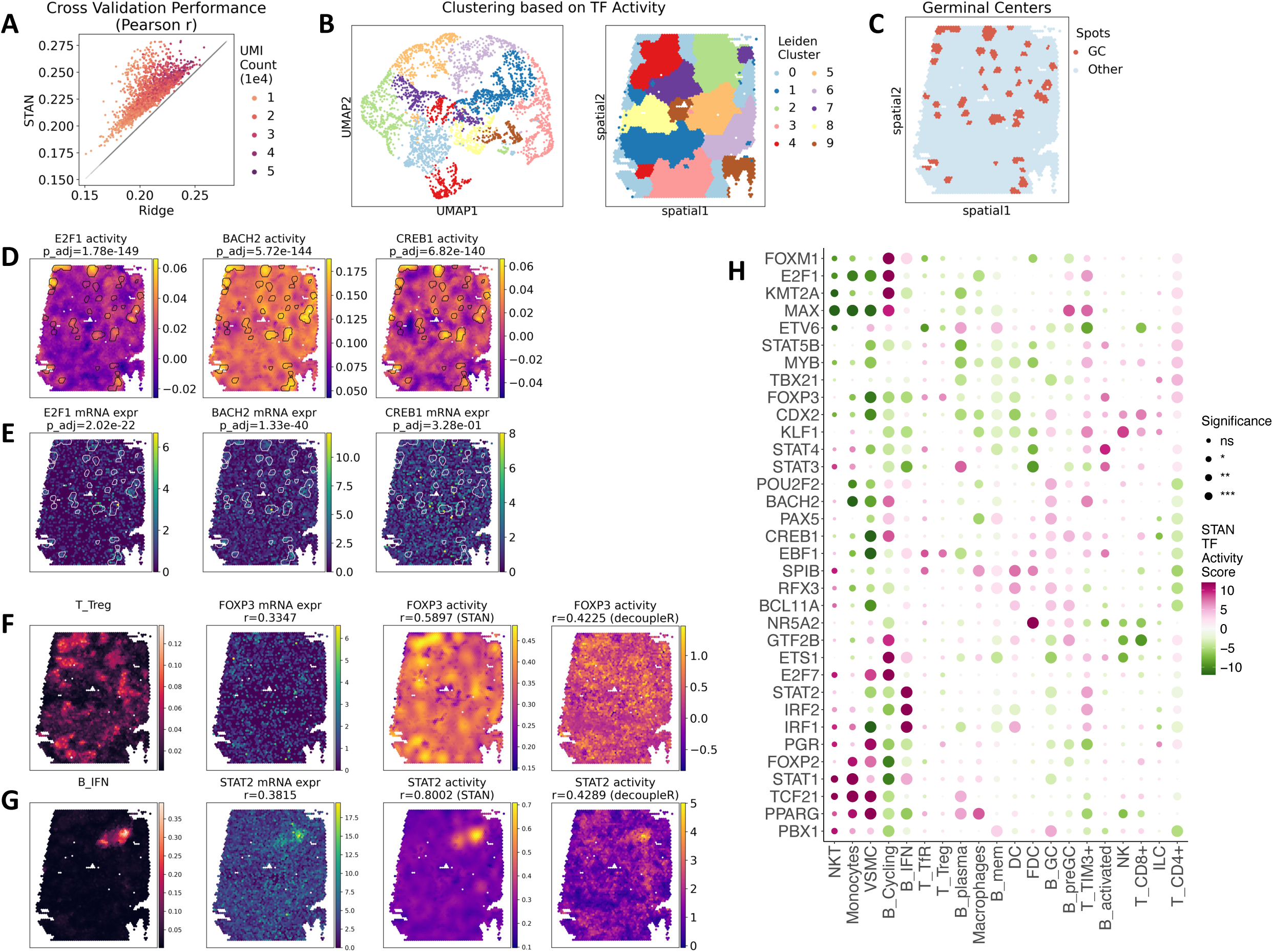
STAN identifies GCs and cell type-specific TFs in a human lymph node. **(A)** STAN and ridge regression models (without spatial information) predict spot-specific expression of held-out genes. Pearson correlations between predicted and actual gene expression for all spots are shown. For each method and each sample, the correlation was computed via 10-fold cross-validation of held-out genes. Each point represents one spot in the dataset and is colored on the basis of the spot’s UMI count. STAN outperforms Ridge for all the spots (dots) above the gray diagonal line. **(B)** UMAP (left) and spatial plot (right) of spots clustered on the basis of inferred TF activity via the Leiden algorithm. **(C)** Locations of GCs. **(D)** Inferred TF activity of the top 3 ranked TFs in the GCs (outlined). **(E)** mRNA levels of the top 3 ranked TFs in GCs (outlined). **(F)** Spatial maps of spot-level proportions of regulatory T cells (left), expression of *FOXP3* (2nd column), inferred activity of FOXP3 via STAN (3rd column), and inferred activity of STAT2 via decoupleR (right). **(G)** Spatial maps of the spot-level proportions of IFN-β cells (left), expression of *STAT2* (2nd column), inferred activity of STAT2 via STAN (3rd column), and inferred activity of STAT2 via decoupleR (right). **(H)** Median TF activity scores across different cell types/states are shown. A TF-cell type pair was selected if the TF activity score was >4, the adjusted *P* value was <1e-2, and r^2^ was > 0.4 for the cell type.

Clustering of the STAN-predicted TF activities identified ten major clusters and revealed that the corresponding spatial distributions presented highly distinct features (**Figure 2B**). The spatial distribution of mRNAs correlated well with TF activity clusters but appeared to be noisier and less precise (**Supplementary Figure 1B**). For example, human lymph nodes are characterized by dynamic microenvironments with many spatially interlaced cell populations, such as GCs. Histological examination of the Visium human lymph node sample revealed multiple GCs [35] (**Figure 2C**); more than half of the annotated GC regions were part of Cluster 0 (**Figure 2B**; **Supplementary Table 4**). We next assessed TF‒GC associations by comparing TF activities between spots in a given GC region and those in all other regions. We identified several known TFs associated with GCs in the lymph node sample (**Figure 2D**), including BACH2, which is critical for the formation and survival of GCs [40], CREB1 [41], and E2F1 [42]. We found fewer associations when TF mRNA expression levels were used directly (**Figure 2E**) or when decoupleR was used [43] without leveraging spatial and morphological features (**Supplementary Figure 1C**).

To identify cell-type-specific TFs, we next used cell type proportions obtained from the Cell2location deconvolution approach [35]. Briefly, Kleshchevnikov et al. analyzed the Visium dataset of the human lymph node and spatially mapped a comprehensive atlas of reference cell types derived by integration of scRNA-seq datasets from human secondary lymphoid organs. To explore the associations between TFs and cell types, we first computed correlations between spot-specific TF activities and cell type proportions. For several known cell type-specific TFs, we observed strong correlations. For example, FOXP3 activity was highly correlated with the regulatory T (Treg) cell type proportion (PCC = 0.5904) (**Figure 2F**), whereas STAT2 activity was highly correlated with the number of cells expressing interferon-β (IFN-β) (PCC = 0.8) (**Figure 2G**). Indeed, FOXP3 is a master TF that regulates Treg cell development and function [44], and STAT2 is essential for immune responses involving Type 1 IFNs, including IFN-β [45]. The correlations between FOXP3/STAT2 activity and the cell type proportion were greater than those between *FOXP3/STAT2* mRNA expression and the cell type proportion (PCC = 0.3223 and 0.3421, respectively; **Figure 2F–G**), highlighting the utility of our method for detecting biologically meaningful patterns in the ST data. In addition, correlations between FOXP3/STAT2 activity and the cell type proportion are greater with inference from STAN than from decoupleR (PCC = 0.4428 and 0.4289, respectively; **Figure 2F–G**), indicating the advantage of incorporating spatial and morphological features in TFA inference.

To systematically identify cell type-specific TFs, we next modeled the relationship between estimated cell type proportions and TF activities via linear regression. This model learns a coefficient for each cell type and for each TF, which we interpreted as the TF score for each cell type across spots. We identified several novel and known TF-cell type relationships in the lymph node sample (**Figure 2H**). These inferred TFs are known regulators of several lymph node cell types, including FOXP3 [44] for Tregs; ETS1 for NK cells [46]; SPI1 (PU1) [47, 48] for macrophages and dendritic cells; BACH2 [49] for B and T cells; and IRF1, IRF2, and STAT2 [50] [45] for TIM^+^ T cells and cells expressing IFN-β. We also identified TFs involved in the control of cell cycle progression, such as MAX, KMT2A, E2F, GTF2B, and FOXM1, that are specifically associated with proliferating B cells but not other B-cell states or other nonproliferative cell types. Importantly, these known relationships were not identified by TF mRNA levels alone (**Supplementary Figure 2A**) or by decoupleR identification of cell type-specific TF activities using only single-cell gene expression measurements (**Supplementary Figure 2B**).

### TF activity patterns across distinct pathological regions in breast cancer

To demonstrate its applicability for analyzing complex cancer tissues, we tested STAN on a publicly available breast cancer ST dataset (n=6) [18]. Leveraging pathological annotations (**Figure 3A** and **Supplementary Figure 3A-B**) from both triple-negative breast cancer (TNBC, n=4) and estrogen receptor-positive breast cancer (ER+, n=2) samples, STAN revealed TF activity patterns across distinct pathological regions in these types of cancer [18] (**Figure 3B** and **Supplementary Figure 4**). We first compared TF activity patterns associated with pathological regions across all breast cancer samples [18], three of which appeared in at least four of the six samples: stroma, lymphocytes, and the combination of invasive cancer, stroma, and lymphocytes (**Figure 3B–C**). We observed that the activity patterns of the TFs associated with the lymphocyte and stroma regions in all the samples were consistent. For the combined invasive cancer, stroma, and lymphocyte regions, we observed heterogeneous TF activity profiles between the TNBC and ER+ samples.

**Figure 3:**
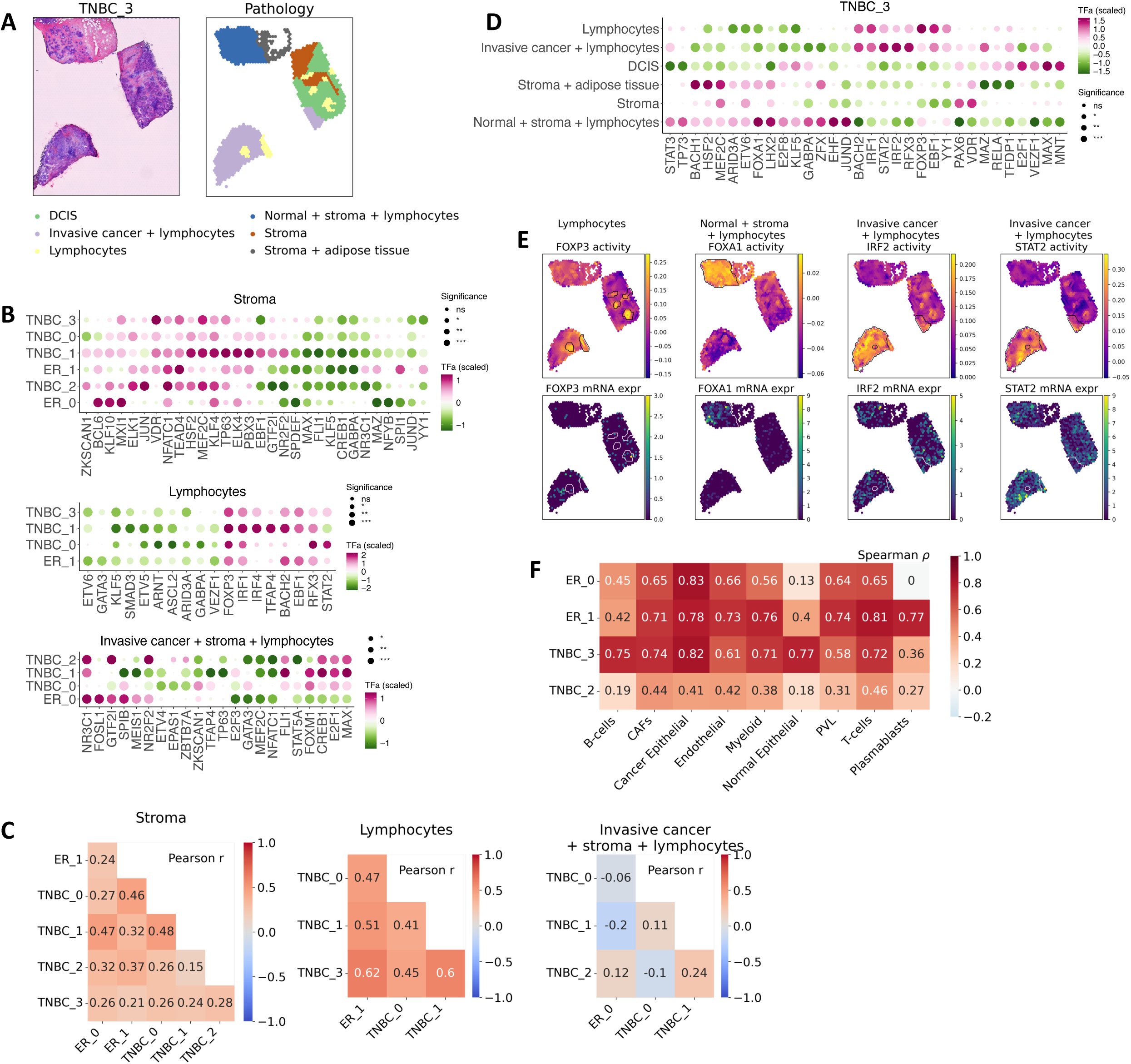
TF activity patterns across distinct pathological regions in breast cancer. **(A)** H&E-stained image (left) and pathological annotation (right) of a triple-negative breast cancer sample from Wu et al. [18] (Sample ID: TNBC_3). **(B)** The mean inferred TF activity differences between a given pathological region [stromal regions (top), lymphocyte regions (middle), and invasive cancer + stroma + lymphocyte regions (bottom)] and all other regions across samples. The dot size indicates −log_10_(FDR) according to the Wilcoxon rank-sum test. **(C)** Heatmaps showing Pearson correlation coefficients between mean TF activities in all applicable samples for stroma (top), lymphocytes (bottom left), and invasive cancer + stroma + lymphocytes (bottom right). **(D)** The mean inferred TF activity differences between a given pathological region and all other regions are shown. The dot size indicates −log_10_(FDR) according to the Wilcoxon rank-sum test. The rows and columns are hierarchically clustered. The list of TFs includes the top 5 TFs ranked by the *P* value associated with each pathological region. **(E)** Spatial feature plots showing FOXP3, FOXA1, IRF2, and STAT2 TF activities (top) and mRNA levels (bottom). **(F)** Correlation between cell type-specific scores based on ST (inferred by STAN) and scRNA-seq (inferred by ridge regression).

In **Figure 3D**, we show TFs associated with different pathological regions for one TNBC sample as a representative relationship. In lymphocyte-rich regions with and without invasive cancer regions, we observed increased activity of IRF, STAT family TFs, FOXP3 (**Figure 3E**), and EBF1. These TFs are important regulators of B and T-cell function and differentiation. We also found that FOXA1 activity was high in invasive cancer regions, the stroma, and lymphocytes (**Figure 3E**). FOXA1 plays pivotal roles in breast development, is crucial for mammary morphogenesis (21,23), mediates full ER activity (24), and directly interacts with the TF GATA3. Notably, the corresponding TF gene expression patterns did not typically match TF activity, except for some TFs whose encoding genes were more highly expressed, such as FOXA1 (**Figure 3E**).

Four ST samples obtained from Wu et al. [18] included scRNA-seq data from the same patient. We inferred cell type-specific TF activities by fitting the model to the scRNA-seq data without spatial information, computed cell type-specific TF scores, and calculated Spearman’s rank correlation coefficient for the TF scores for each cell type between the corresponding ST and scRNA-seq data (**Figure 3F**). The medium-to-high correlation for most cell types and samples was consistent between the scRNA-seq data and the ST-based inference, although the ST data were sparser.

### Linking ligands and receptors to TFs in glioblastoma

Cell behavior is intricately influenced by signals from the surrounding microenvironment, often in the form of ligands, which are extracellular protein signals produced by neighboring cells. These ligands interact with receptors expressed on recipient cells, triggering a cascade of molecular events that can profoundly impact gene expression programs and modulate the activity of TFs. Complexity and heterogeneity have been demonstrated to significantly contribute to tumor aggressiveness and poor prognosis in glioblastoma (GBM), the most common and deadliest type of brain cancer. A wide variety of nonneoplastic stromal cells, extracellular matrix (ECM), and tumor cells are present in anatomically distinct regions to form various tumor niches, which profoundly influences the complexity of GBM [51]. To further illustrate these microenvironmental niches by exploring the molecular interplay between ligands, receptors, and TFs, we leveraged STAN to analyze Visium ST data obtained from one human glioblastoma sample (**Supplementary Figure 5**). We employed inferred TF activities to model intercellular signaling pathways originating from ligands expressed within the local neighborhood of cellular regions and to explore the complex communication networks within the tumor microenvironment.

We began by determining the average normalized mRNA counts of neighboring spots for a selection of ligands extracted from the CellTalkDB database [52], which were individually calculated for each spot. We subsequently examined the correlation between the mean mRNA levels of neighboring spots and the inferred TF activity and identified patterns in the interplay between ligand and receptor expression, TF activity, and the cellular microenvironment (**Figure 4A**) through two-way clustering of ligands, receptors, and TFs on the basis of pairwise PCCs. For example, our analysis revealed a significant correlation (PCC > 0.6) between STAN-predicted SOX2 and HIF1A activity and the expression levels of RACK1, CD44 receptors, VIM (vimentin), VEGFA, and DKK1 ligands (**Figure 4A-B**). SOX2 has been extensively implicated in promoting malignancy in glioblastoma, with its regulatory roles extending to CD44, DKK1, and VIM, among other factors [53]. Additionally, RACK1, acting as a scaffolding protein, modulates the expression of pluripotency TFs such as Nanog, Oct4, and SOX2 in human hepatocellular carcinoma [54].

**Figure 4:**
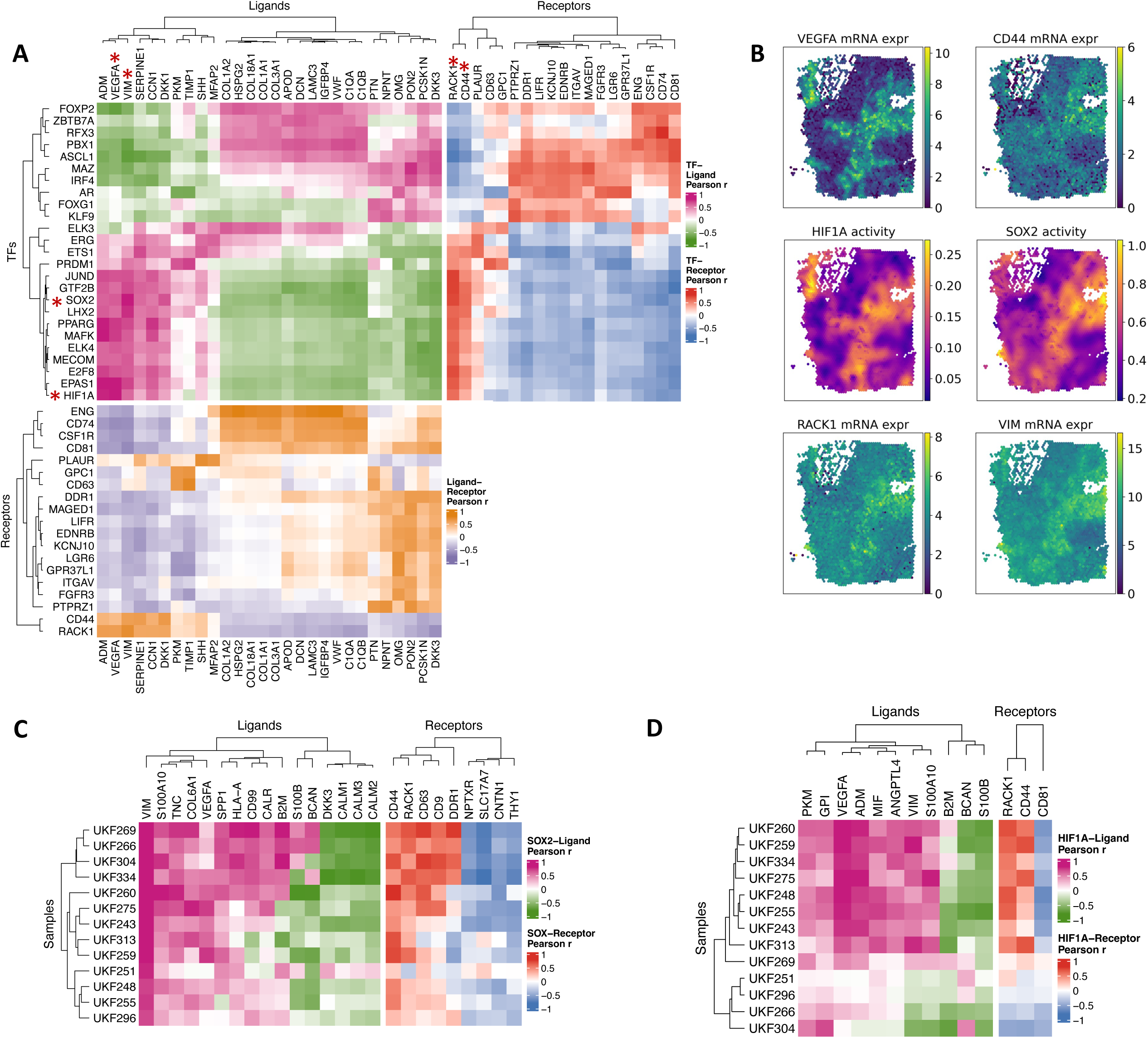
STAN links ligands/receptors to TFs in glioblastoma. **(A)** Clustered heatmap showing the Pearson correlation coefficient between inferred TF activities and ligand/receptor expression of the 10x Genomics Human Glioblastoma sample. **(B)** Spatial feature plots displaying the mRNA expression of *VEGFA* and *CD44* (ligands) (top), inferred activity of *HIF1A* and *SOX2* (middle), and mRNA expression of *RACK1* and *VIM* (receptors) (bottom). (C, D) Clustered heatmaps showing the Pearson correlation coefficients between inferred SOX2 and HIF1A activities and ligand/receptor expression in thirteen glioblastoma samples from Ravi et al.[55].

To extend our analysis of VEGFA/RACK1-HIF1A and VIM/CD44-SOX2, we applied STAN to an additional thirteen spatial tracsriptomics GBM samples from Ravi et al. [55]. We correlated the inferred activity of SOX2 and HIF1A with the expression levels of their respective ligands and receptors. We selected ligands and receptors with a significant correlation (PCC > 0.5) in at least five samples (**Figure 4C**). Our analysis revealed that SOX2 activity was highly correlated with the expression of CD44 and VIM across all samples. Similarly, HIF1A activity showed a strong correlation with the expression of RACK1 and VEGFA, particularly in samples with medium or low proportions of neuronal structures [56] (**Supplementary Fig. 6**).

To further explore the coupling of SOX2 with CD44 and VIM and the coupling of HIF1A with RACK1 and VEGFA, we performed IF on glioblastoma patient samples. Indeed, we detected the coexpression of SOX2, CD44, and VIM as well as HIF1A, RACK1 and VEGFA at the protein level (**Figure 5A-B**). In addition, we compared immunohistochemically (IHC)-stained images of ligands, receptors and TFs in glioblastoma tissues obtained from the Human Protein Atlas (HPA) database [57]. IHC images revealed that the TFs ZBTB7A, PPARG, MECOM, JUND, PRDM1, HIF1A, SOX2, FOXP2 and GTF2B and the ligands and receptors HSPG2, VEGFA, ADM, GPC1, CD81, and VIM were expressed in glioblastoma tissues (**Supplementary Fig. 7**). These results indicate that our computational approach, combined with experimental validation, can elucidate the landscape of the tumor microenvironment on the basis of molecular communication within ligand–receptor–TF networks.

**Figure 5:**
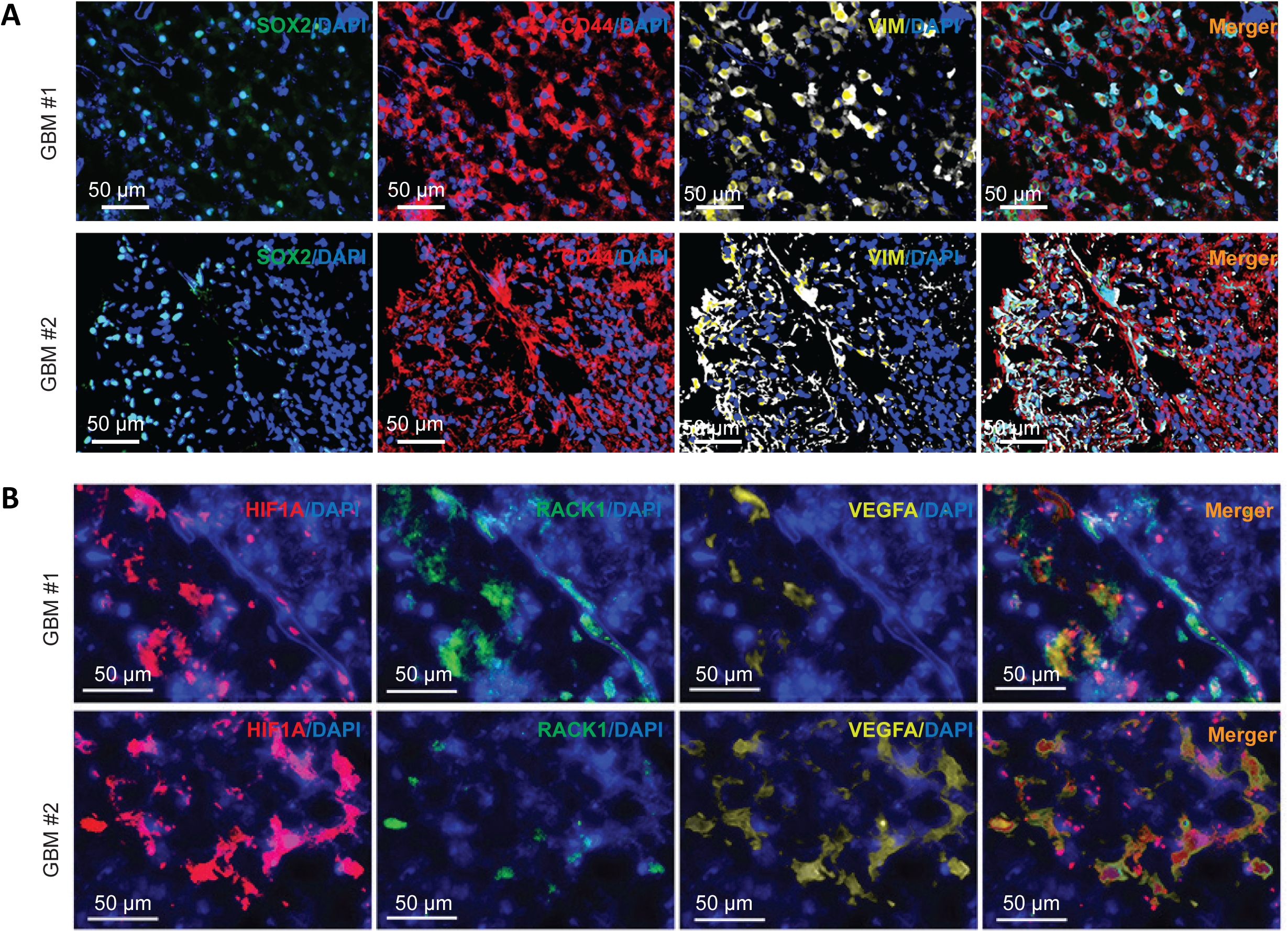
Representative images of glioblastoma patient tumors costained for **(A)** SOX2, CD44, and VIM, and **(B)** HIF1A, RACK1, and VEGFA.

## Discussion

ST datasets provide rich information about cellular diversity and heterogeneity, as well as the dynamics of complex biological systems. One advantage of ST data is the additional information of co-registered images, which contain both morphological and functional patterns and distance information. Here, we present STAN, a broadly applicable tool that leverages parallel spot-specific ST data and imaging data with *cis*-regulatory information (e.g., TF-target gene priors) to predict spatially informed TF activities. While several methods for inferring TF activities use TF expression to predict gene expression, they do not consider how cell identity is influenced by neighboring cells.

We utilized STAN to uncover TF relationships across diverse tissue types, from lymph nodes to more complex tissues, such as breast cancer and glioblastoma. Applying STAN to ST datasets enabled us to (i) decipher critical TF regulators underlying cell identities, spatial domains (e.g., GCs), and pathological regions (e.g., stroma vs. tumor); (ii) determine whether a given pathological region/spatial domain has different and/or common regulators across disease subtypes (e.g., stroma in TNBC vs. ER+ breast cancer); (iii) identify similar and/or different TFs associated with spatial domains or cell types across healthy individuals and those manifesting a disease; and (iv) link ligands and receptors to TFs to elucidate potential signaling pathways and regulatory networks involved in cellular communication and tissue microenvironment interactions. We released STAN as an open-source software package to facilitate further studies and enable researchers to quantify the activity of important transcription factors that are normally obscured by their modest mRNA-level expression.

The identified TFs from our analysis of lymph node, glioblastoma, and breast cancer data are clinically and biologically relevant. The distinctive activity patterns of these TFs may offer valuable insights for diagnostic and prognostic purposes in the management of breast cancer, glioblastoma, and other cancer types. Within a tumor, several subpopulations of cancer cells can differ from each other in their context-specific TFs, which drive their functions because they reside in distinct tumor microenvironments. Accurately identifying context-specific TFs, such as those associated with malignant cells in regions with a greater degree of infiltrating immune cells than those without, is crucial. Furthermore, we used STAN to delineate TF, ligand and receptor relationships. For example, our analysis revealed a significant correlation (PCC > 0.6) between STAN-predicted SOX2 activity and the expression levels of CD44 receptors and the VIM (vimentin) ligand. We validated the coexpression of SOX2, VIM and CD44 at the protein level in independent glioblastoma patient samples. This validation suggests promising biomedical applications, such as targeted therapies, and encourages further exploration of these regulators in glioblastoma. Future validation experiments can evaluate the role of SOX2 coupling with VIM and CD44 in glioblastoma. SOX2 expression can be evaluated in the context of VIM and CD44 modulation in cultured glioblastoma cells. Ultimately, in vivo validation of the role of SOX2, VIM and CD44 can be performed via glioblastoma mouse models through the silencing of SOX2 and/or VIM and CD44 and the measurement of the functional activity of glioblastoma cells. Overall, STAN is applicable to any disease type (e.g., autoimmune diseases) or biological system (e.g., immune system) and will enable researchers to leverage spot-based ST datasets to predict context-specific TF activities.

The method we describe has several limitations. First, measurements from capture locations (spots) often include mixtures of multiple cells, leading to analytical challenges in dissecting cellular disposition, particularly in complex systems, such as cancerous tissues. However, the STAN framework can be extended to model emerging single-cell resolution STs, as we will report elsewhere. Second, STAN relies on curated TF target–gene interactions from various sources (e.g., literature-curated data, ChIP–seq peaks, and TF binding site motifs) that are often noisy, incomplete, and not context specific. Our framework can be extended by emerging spatial epigenome‒transcriptome coprofiling [58] for more accurate representation of both promoter and enhancer regions, as performed in the context of patient-specific predictive regulatory models [59, 60]. In addition, STAN does not account for directionality in the TF-target gene interaction matrix. Thus, negative values of inferred TF activities should be interpreted on the basis of whether the TF acts as an activator or repressor. Our model assumes that a TF either induces or represses its target(s), but some TFs may switch roles on the basis of cofactors, potentially confounding data interpretation. Finally, we use a fixed gene‒target representation, inferring TF activity by correlating it with target expression via a linear model, which does not capture the more complex combinatorics of TF binding.

## Conclusions

In summary, STAN will enable more robust and context-specific analysis of cellular regulatory states in heterogeneous and complex tissue types based ST datasets.

## Methods

### Data and preprocessing

We obtained three publicly available genome-wide ST datasets comprising one human lymph node sample and one glioblastoma sample (both from the 10x Genomics website), along with six breast cancer samples derived from multiple patients from Wu et al. [18] and thirteen GBM data derived from multiple patients from Ravi et al. [55] (**Supplementary Table 1**). These datasets represent both structured (e.g., lymph node) and unstructured (e.g., solid tumors) tissues. For each ST dataset, we collected spot-level expression data, spatial coordinates of the spots, and the associated image of the tissue slice. Four ST samples obtained from Wu et al. [18] included scRNA-seq data from the same patient. We also downloaded these processed scRNA-seq data with cell type annotations [18] (GSE176078).

We obtained predicted cell type abundance scores and GC annotations for the lymph node data from Kleshchevnikov et al. [35] and pathological annotations for the breast cancer dataset from Wu et al. [18]. We obtained a gene set resource comprising TF-target gene priors from hTFtarget [61] and retained only those TFs identified in the Human Transcription Factor database [62] to generate the TF-target gene prior matrix, which delineates a candidate set of associations between TFs and target genes.

We then analyzed the ST data via Scanpy (version 1.9.3) [63] and anndata (version 0.9.2) [64]. Genes expressed in fewer than five spots were filtered, and low-quality spots were filtered on the basis of unique molecular identifier (UMI) counts (dataset-specific thresholds, **Supplementary Table 2**). For each ST dataset, the TF-target gene prior matrix (referred to as **D**) and spot-level gene expression matrix (referred to as **Y**) underwent a sequential filtering process. Initially, genes expressed in less than 20% of the filtered spots were eliminated. Only the mutual genes present in both matrices were subsequently retained. Further refinement involved removing genes associated with fewer than five remaining TFs. Finally, TFs associated with fewer than 10 remaining genes were removed to address insufficient target gene representation for reliable analysis. For each filtered gene expression matrix, we normalized each spot by total counts over all genes in that dataset. The normalized counts were then square root transformed to stabilize the variance.

### Training sample-specific STAN models

STAN is designed to leverage *cis*-regulatory information (e.g., TF-target gene priors), gene expression, spatial information, and morphological features extracted from corresponding H&E staining imaging data acquired via ST to enable inference of spatially informed, spot-specific TF activities. **Figure 1** illustrates our overall STAN framework. We let **D**ɛR^(*N*×*Q*)^ and **Y**ɛR*^N^*^×*M*^ denote the TF-target gene prior matrix and the normalized spot-level gene expression matrix, where *N*, *M*, and *Q* represent the numbers of genes, spots, and TFs, respectively. We subsequently formulate a regression problem aimed at learning the weight matrix **W**ɛR^(*Q*×*M*)^ between TFs and spots to predict TF-target gene expression through the following equation:

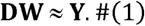

To include spatial features in Equation (1), we define a Gaussian kernel matrix **K**ɛR*^M^*^×*M*^ as follows. For the *i*th spot, we first cropped the image to a square of size (2*a* + 1) centered at the spot and then calculated the pixel intensity of the spot as the mean RGB value across all the pixels in the cropped image. The spatial coordinate of the spot, (*x_i_*, *y_i_*) as well as the RGB pixel intensity of the spot (*r_i_*, *g_i_*, *b_i_*), forms a 5-dimensional vector v^(*i*)^ = (*x_i_*, *y_i_*, *ωr_i_*, *ωg_i_*, *ωb_i_*), which contains both the spatial and morphological information of the *i*th spot [65], and ω is a weight parameter multiplied by its morphological components. Next, we normalized each component in v to have a zero mean and unit variance across all the spots. We let *b* be the parameter that determines the bandwidth of the Gaussian kernel. Given two spots indexed *i* and *j*, the(*i,j*)th element of K is given by:

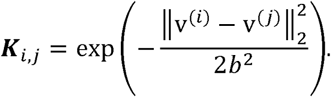

Note that the kernel K is symmetric (i.e., **K** = **K***^T^*).

We assume that the TF activity matrix **W** can be split into two matrices:

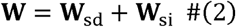

where **W**_sd_ is spatially and morphologically dependent and where **W**_si_ is spatially and morphologically independent. With the kernel **K** defined, to find the optimal solution **W**\* of Equation (1), which has the minimal residual while reserving as many spatial features as possible, we solve Equation (1) via spatially weighted ridge regression:

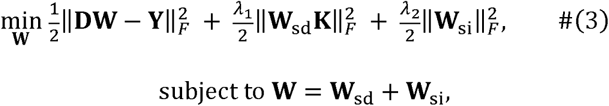

where *F* denotes the Frobenius norm and where λ_1_, λ_2_ are positive hyperparameters. The nonspatial component **W**_si_ is *L*_2_-regularized, whereas the *L*_2_-regularizer of the spatial component **W**_sd_ is weighted by the kernel **K**. Equation (3) can be solved efficiently via a low-rank approximation of the spanning set of **K**.

The analytic solution of STAN can be derived as follows. Let *F* be the objective function of the optimization problem (3), that is:

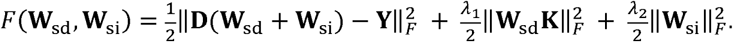

The gradients of *F* with respect to **W**_sd_ and **W**_si_ are as follows:

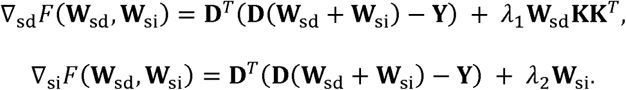

The optimal solution 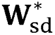 and 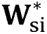 that minimize *F* satisfy 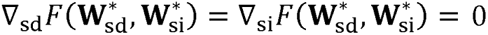 are as follows:

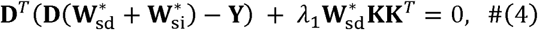

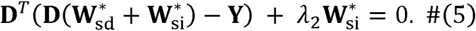

Comparing Equations (4) and (5), we can observe the following:

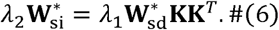

Combining Equations (2) and (6), the optimal solution **W**\* of Equation (1) can be simplified as:

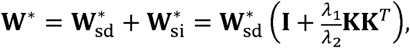

where **I** is the identity matrix. Thus, it is sufficient to solve 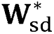.

From Equation (4),

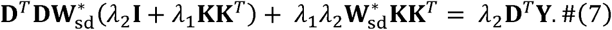

Reformulating the matrix products in Equation (7) yields an equivalent linear system:

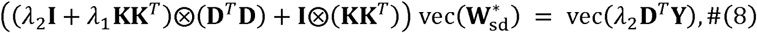

where 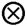 is the Kronecker multiplication and where *vec* is the vectorization of a matrix, we make use of the symmetric properties of **I** and **K**. From Equation (8), we have:

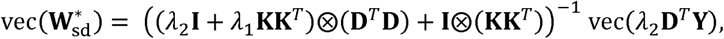

which can be solved efficiently via singular value decomposition (SVD).

For comparison, we also solve Equation (1) via ridge regression:

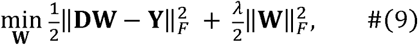

which does not model any spatial features, and the solution from Equation (9) was considered the baseline. We evaluated performance with 10-fold cross-validation and computed the Pearson correlation coefficient (PCC) between the predicted and measured gene expression profiles at held-out spots. Finally, we use the trained **W** to infer the TF activities for each spot. To cluster gene expression and STAN-predicted TF activities (**W**, scaled to unit variance and zero mean), we employed the Leiden algorithm (51). Additionally, a neighbor graph of spots was generated and embedded via uniform manifold approximation and projection (UMAP) (49,50).

### Deconvoluting ST data

We employed GraphST (version 1.1.1) [66] to perform cell-type deconvolution on human breast cancer samples, integrating both the scRNA-seq and the ST data [18]. GraphST is a graph-based self-supervised contrastive learning method that makes use of both spatial information and gene expression profiles for spatially informed clustering, batch integration, and cell-type deconvolution. By utilizing the shared samples (**Supplementary Table 3**), we applied the GraphST deconvolution pipeline with the recommended settings. Following training of the graphical model, we derived a mapping matrix wherein each element represents the mapping probability of a cell type in each spot.

### Inferring cell type-specific TFs

Widely used ST platforms, such as Visium, measure the whole transcriptome in low-resolution spots that may capture multiple cells. To identify cell type-specific TFs via STAN, we conducted association analysis between TFs and cell types by predicting the activity of each TF individually from spot-specific cell type proportions. These proportions were estimated via deconvolution methods (e.g., Cell2location [35], GraphST [66], DestVI [36], Tangram [37], Stereoscope [38], and RCTD [39]) to integrate spatial RNA-seq data with reference cell-type signatures obtained from scRNA-seq profiles of the same tissue type. Briefly, we let **A** denote each spot-specific cell type matrix, where each row of **A** represents a spot and each column a cell type. For the *i*th TF, we let *w_i_* be a column of **W**, which is the inferred activity of the TF across all spots. Then, by using a linear regression model (the ordinary least squares method), we inferred the cell type-specific score *x_i_* of the TF via **A***x_i_* = *w_i_*, where each component of *x_ij_* is the TF score for the *j*th cell type. We evaluated the significance of the effect size (coefficient) for each cell type in the regression model via Student’s t distribution under the null hypothesis. Following false discovery rate (FDR) correction across TFs, we identified a significant set of TFs whose activities are associated with each cell type.

### Identifying pathological region-specific TFs

To identify pathological region-specific TFs, we initially scaled the inferred TF activity matrix to unit variance and zero mean. For each region and each TF, we subsequently conducted a Wilcoxon rank-sum test to compare the inferred TF activity within and outside the region. Finally, we adjusted the *P* values via the Benjamini‒Hochberg (BH) procedure.

### Linking TFs to ligands and receptors

We obtained a list of human ligand‒receptor interaction pairs from the CellTalkDB database [52], which includes 3398 available ligand‒receptor interaction pairs from among 815 ligands and 780 receptors. We began by determining the average normalized mRNA counts of neighboring spots for the ligands and receptors individually calculated for each spot. We subsequently computed the PCC between the mean mRNA levels of neighboring spots and the inferred TF activities. Highly correlated TF‒ligand, TF‒receptor, and ligand‒receptor pairs were selected on the basis of thresholds (PCC > 0.6), and PCCs were visualized in clustered heatmaps.

### Implementing decoupleR

We compared STAN with decoupleR (version 1.5.0) [43], which can be used to extract biological activities (e.g., transcription factor activities) from expression-level data. At the recommended settings for inferring TF activity, we ran decoupleR for the filtered human lymph node sample (**Supplementary Table 2**).

### Human protein atlas

The Human Protein Atlas (https://www.proteinatlas.org) is a public resource that extracts information, including images from immunohistochemistry (IHC), protein profiling, and pathologic information, from specimens and clinical material from cancer patients to determine global protein expression [67]. Here, we compared the protein expression of available TFs, ligands and receptors in glioblastoma tissues via IHC images.

### Immunofluorescence (IF)

For IF staining, OCT-embedded frozen brain tumor sections were thawed at room temperature for 30 min, rinsed and rehydrated with TBS 3 times, and the tissue was fixed for 10 min in acetone. After being blocked with normal horse serum (2.5%), the sections were incubated with the indicated primary antibodies overnight at 4°C. The specific dilutions of the primary antibodies used were as follows: rabbit SOX2 (1:200; EMD Millipore AB5603), rat CD44 (1:100; Biolegend 103002), mouse VIM (1:100; Biolegend 677801), rabbit HIF1A (1:100; Abcam ab308433), goat RACK1 (1:30; R&D AF3434), and mouse VEGFA (1:200; Abcam ab1316). After three washes with TBST (0.1% Tween), the slices were incubated with goat anti-rabbit IgG H&L Alexa Fluor 488 (1:500; Invitrogen A11008), donkey anti-rat IgG H&L Alexa Fluor 594 (1:500; Invitrogen A21209) and goat anti-mouse IgG H&L Alexa Fluor 647 (1:500; Invitrogen A32728) at RT for 1 h. After being washed with TBST three times, the slides were mounted with DAPI (Abcam ab104139) and imaged via a Leica DMi8 confocal microscope. Frozen glioblastoma (GBM) samples from The Cancer Genome Atlas (TCGA) were collected as described previously [68].

### Statistical analysis and visualization

Statistical analysis and visualization were performed in Python (version 3.11.5) via the SciPy (version 1.10.1) and statsmodels (version 0.14.0) packages. For general data analysis and manipulation, pandas (version 2.0.3) and NumPy (version 1.22.4) were used. Graphs were generated via the Python packages matplotlib (version 3.7.2) and seaborn (version 0.12.2) and the R (version 4.4.0) packages ggplot2 (version 3.5.1) and ComplexHeatmap (version 2.20.0).

For comparisons of inferred TF activities between groups, we performed a Wilcoxon rank-sum test and determined the direction of shifts by comparing the means of the two populations. We corrected the raw P values for multiple hypothesis testing via the Bonferroni and FDR (BH) methods.

## Declarations

### Data availability

The 10x Genomics datasets were downloaded via the Scanpy function scanpy.datasets.visium_sge with sample IDs V1_Human_Lymph_Node and Parent_Visium_Human_Glioblastoma. The cell type and geminal center annotations were downloaded from the Cell2 location study by Kleshchevnikov et al. [35] (https://github.com/vitkl/cell2location_paper/tree/master/notebooks/selected_results/lymph_nodes_analysis). The ST breast cancer data from Wu et al. [18], including pathological annotations, were accessed through the Zenodo data repository (https://doi.org/10.5281/zenodo.4739739). The ST glioblastoma data from Ravi et al. [55], including the spatial metaprogram annotations, were accessed through the GitHub repository (https://github.com/tiroshlab/Spatial_Glioma) provided by Greenwald et al [56]. The scRNA-seq data from Wu et al. [18] were obtained from the Gene Expression Omnibus (GEO) dataset under accession number GSE176078. A list of human ligand‒receptor interaction pairs was obtained from the CellTalkDB database [52] (https://github.com/ZJUFanLab/CellTalkDB/blob/master/database/human_lr_pair.rds).

### Code availability

A Python package code for STAN is available at https://github.com/osmanbeyoglulab/STAN. The codes used to produce the results in this paper, including those used for data preprocessing, cell type deconvolution, model fitting, and downstream analyses, are available as Jupyter notebooks in the same GitHub repository.

## Ethics approval and consent to participate

The study analyzed publicly available sequencing data.

## Consent for publication

Not applicable.

## Competing interests

The authors declare that they have no competing interests.

## Authors’ contributions

LZ performed computational experiments and analyses, helped to develop the algorithmic approaches and helped to write the paper. AS helped to develop the algorithmic approaches. QB performed the experimental validation and helped to write the experimental validation section. ESK gathered IHC images for the validation. BH supervised the experimental validation. HUO supervised the project, helped to develop the algorithmic approaches and helped to write the paper.

## Acknowledgments

We thank Jing Hong Wang, Anthony R. Cillo, and Harinder Singh for helpful discussions. HUO was supported by the National Institutes of Health (R35GM146989). LZ was supported by the National Natural Science Foundation of China (12101342) and the Ningbo Yongjiang Talent Introduction Programme (2022A-222-G). Data analyses were supported by the University of Pittsburgh Center for Research Computing and the Extreme Science and Engineering Discovery Environment (XSEDE) supported by the National Science Foundation (OCI-1053575) via the Bridges2 system supported by the National Science Foundation (ACI-1445606) at the Pittsburgh Supercomputing Center.

## Notes

### Competing Interest Statement

The authors have declared no competing interest.

### Summary of Updates

Added one author and one figure

